# The C-type Lectin Receptor Dectin-2 is a receptor for *Aspergillus fumigatus* galactomannan

**DOI:** 10.1101/2022.04.12.488040

**Authors:** Jennifer L. Reedy, Arianne J. Crossen, Paige E. Negoro, Hannah E. Brown, Rebecca A. Ward, Diego A. Vargas Blanco, Kyle D. Timmer, Michael K. Mansour, Marcel Wüthrich, Thierry Fontaine, Jean-Paul Latgé, Jatin M. Vyas

## Abstract

*Aspergillus fumigatus* is a ubiquitous environmental mold that causes significant mortality particularly amongst immunocompromised patients. The detection of the *Aspergillus-*derived carbohydrate galactomannan in patient sera and bronchoalveolar lavage fluid is the major biomarker used to detect *A. fumigatus* infection in clinical medicine. Despite the clinical relevance of this carbohydrate, we lack a fundamental understanding of how galactomannan is recognized by the immune system and its consequences. Galactomannan is composed of a linear mannan backbone with galactofuranose sidechains and is found both attached to the cell surface of *Aspergillus* and as a soluble carbohydrate in the extracellular milieu. In this study, we utilized fungal-like particles composed of highly purified *Aspergillus* galactomannan to identify a C-type lectin host receptor for this fungal carbohydrate. We identified a novel and specific interaction between *Aspergillus* galactomannan and the C-type lectin receptor Dectin-2. We demonstrate that galactomannan bound to Dectin-2 and induced Dectin-2 dependent signaling including activation of spleen tyrosine kinase and potent TNFα production. Deficiency of Dectin-2 increased immune cell recruitment to the lungs but was dispensable for survival in a mouse model of pulmonary aspergillosis. Our results identify a novel interaction between galactomannan and Dectin-2 and demonstrate that Dectin-2 is a receptor for galactomannan which leads to a pro-inflammatory immune response in the lung.

**IMPORTANCE:** *Aspergillus fumigatus* is a fungal pathogen that causes serious and often fatal disease in humans. The surface of *Aspergillus* is composed of complex sugar molecules. Recognition of these carbohydrates by immune cells by carbohydrate lectin receptos can lead to clearance of the infection or, in some cases, benefit the fungus by dampening the host response. Galactomannan is a carbohydrate that is part of the cell surface of *Aspergillus* but is also released during infection and is found in patient lungs as well as their bloodstreams. The significance of our research is that we have identified a mammalian immune cell receptor that recognizes, binds, and signals in response to galactomannan. These results enhance our understanding of how this carbohydrate interacts with the immune system at the site of infection and will lead to broader understanding of how release of galactomannan by *Aspergillus* effects the immune response in infected patients.

## INTRODUCTION

*Aspergillus fumigatus* is a saprophytic mold that causes a wide spectrum of clinical manifestations ranging from allergic disease to invasive infections, including pneumonia and disseminated disease. While the majority of patients that develop invasive aspergillosis are immunocompromised due to transplantation, inherited immune deficiencies, or use of immunosuppressant therapies, invasive aspergillosis also occurs in otherwise healthy individuals in association with preceding influenza or SARS-CoV2 infection (1–3). The mortality rate for invasive aspergillosis remains unacceptably high with approximately 65 – 80% of patients succumbing to disease despite antifungal therapy (4), and is a major cause of morbidity, particularly among patients with compromised immune systems.

Humans are infected with *A. fumigatus* through inhalation of conidia deposited within small airways and alveoli. Conidia are eliminated in part by mechanical expulsion through the action of mucus and ciliated cells, as well as killing by macrophages and neutrophils (5, 6). Upon encountering a fungal pathogen, innate immune cells synchronously engage multiple fungal cell wall antigens through a diverse array of pattern recognition receptors including Toll-like receptors, scavenger receptors, complement receptors, and C-type lectin receptors (CLRs), and integrate those signals to generate a coordinated immune response (7–11). The majority of the fungal cell wall antigens are polysaccharides, as > 90% of the fungal cell wall is composed of complex carbohydrates. The cell wall is commonly composed of an inner chitin and middle β-1,3 glucan layer with additional carbohydrate constituents like an outer mannan layer depending upon the genus of fungus (12, 13). In *A. fumigatus*, the mannan content is primarily exists within the galactomannan polymer. Although specific cell wall carbohydrate–pattern recognition receptor interaction pairings have been defined for a few cell wall polysaccharides, such as β-1,3 glucan and the CLR Dectin-1 (8, 14), we lack a fundamental understanding of the specific receptors that engage many other fungal carbohydrates and how these ligand-receptor cognate pairs drive the innate immune response.

The fungal cell wall is a highly dynamic structure that changes in both structure and composition depending on the morphological stage, environmental milieu, and upon engagement with immune cells (13, 15). Therefore, probing the role of a single carbohydrate entity using whole pathogens is challenging. Standard techniques have relied upon solubilized carbohydrates or genetic deletion mutants, however these techniques have limitations including pleiotropic changes that occur in genetic deletion strains, such as compensatory changes in cell wall composition or structure, growth defects, alterations in antigen exposure (16), and known differences in immune response to soluble versus particulate (or cell-bound) carbohydrates. Most fungal cell wall carbohydrates remain bound to the cell wall, however galactomannan and β-1,3 glucan are unique as they exist as both cell wall-associated and free/soluble carbohydrates. β-1,3 glucan has been extensively studied revealing differences in how soluble and cell wall β-1,3 glucan interact with immune cells (17–19). Cell wall associated β-1,3 glucan signals via both the CLR Dectin-1 and complement receptor 3, whereas soluble β-1,3 glucan is Dectin-1 independent (17–19). Although both cell wall β-1,3 glucan and soluble β-1,3 glucan bind Dectin-1, only cell wall β-1,3 glucan activates signaling (17). In contrast, receptors such as TLRs signal in response to both soluble and cell associated ligands. Activation of Dectin-1 requires receptor clustering into phagocytic synapses which exclude regulatory phosphatases licensing activation of Dectin-1 (17, 20). Whether the difference in response to cell wall and soluble carbohydrates is unique to β-1,3 glucan and Dectin-1 or extends to other fungal carbohydrates and CLRs remains unknown. We hypothesize that the innate immune system distinguishes between soluble and cell-associated fungal carbohydrates to tailor the immune response. Most studies to date have been conducted using soluble galactomannan. Thus, we sought to characterize the immune response to cell-associated galactomannan to understand how innate immune cells interact with *Aspergillus* at the site of infection.

The carbohydrate galactomannan was identified approximately 40 years ago as an important clinical biomarker for the diagnosis of invasive aspergillosis, as it can be detected in serum and bronchial fluid from infected patients (21–24). However, the biological and immunological responses to this carbohydrate remain poorly defined. Galactomannan is composed of a linear mannan core composed of α-1,2-linked mannotetraose units attached via a-1,6-linkage with side changes composed of four to five β-1,5-linked galactofuranose residues (25, 26). Galactomannan can be found in both the conidial and mycelial stages of the *Aspergillus* lifecycle. It is bound to the cell membrane through a GPI-anchor, covalently attached to β-1,3-glucan in the cell wall and also released as a soluble polysaccharide into the extracellular milieu (27, 28). Most studies have utilized soluble galactomannan, since strains of *Aspergillus* that lack galactomannan have altered growth morphology and kinetics (29, 30). Stimulation of peripheral blood mononuclear cells (PBMCs) with soluble galactomannan suppressed LPS-induced cytokine production. Furthermore, an *Aspergillus* vaccine protection model demonstrated that treatment with soluble galactomannan induced a non-protective Th2/Th17 immunity (31, 32). Dendritic cell and macrophages internalization of *A. fumigatus* conidia by the C-type Lectin Receptor DC-SIGN could be inhibited through the addition of soluble galactomannan (33), suggesting that DC-SIGN could play a role in recognizing this carbohydrate. These observations indicated that *Aspergillus* subverts the immune response by shedding galactomannan, but the innate immune response to cell-associated galactomannan has yet to be determined.

In this study, we identified Dectin-2 as a receptor for cell wall-associated galactomannan by using fungal-like particles (FLPs), a novel technique previously developed in our laboratory (34). To determine how the immune system recognizes galactomannan in the cell-associated state, we isolated galactomannan from *A. fumigatus* and created FLPs consisting of purified galactomannan. Using these particles to screen a library of CLR reporter cells, we identified Dectin-2 as a receptor for galactomannan. We confirmed the binding of Dectin-2 to galactomannan FLPs, using a solubilized receptor and demonstrated galactomannan FLPs stimulate production of TNFα by murine macrophages in a Dectin-2 dependent manner. Furthermore, we determined that immunocompetent mice lacking Dectin-2 recruit increased numbers of immune cells to lungs compared with wild-type mice in a model of pulmonary aspergillosis, but Dectin-2 is dispensable for survival. Taken together, our results demonstrate a novel and important role for Dectin-2 in innate immune recognition of *Aspergillus* galactomannan.

## RESULTS

### Dectin-2 is a receptor for galactomannan

Since CLRs are known to be involved in sensing of fungal pathogens and bind to carbohydrates, we examined if any of the known CLRs were potential receptors for galactomannan in the *Aspergillus* cell wall. We first screened a murine CLR library using *A. fumigatus* germlings to determine which CLRs were stimulated by the whole organism and confirm utility of the assay. T cell hybridomas containing an NFAT-lacZ reporter were engineered to express individual or combinations of CLRs: Dectin-1, Dectin-2, Dectin-3 (MCL), or Mincle. Dectin-2, Dectin-3, and Mincle require interaction with the gamma subunit of the Fc gamma receptor (FcRγ) for both cell surface localization and signaling, thus this protein was co-expressed. We stimulated the CLR reporter cells for 18 hours with live and heat-killed *Aspergillus* germlings and observed robust activation of Dectin-1, Dectin-2, and Dectin-2/Dectin-3 co-expressing cells (Figure 1A). Since cells that express Dectin-3 alone were not activated, the activation of the Dectin-2/Dectin-3 co-expressing cells is likely due to either activation of Dectin-2 alone or possible Dectin-2/Dectin-3 in combination (Figure 1A). Activation of Dectin-1 is known to occur through binding of cell-wall associated β-1,3 glucan, but the Dectin-2 ligand on the surface of *Aspergillus* has not been established. We hypothesized that galactomannan could be the potential Dectin-2 ligand.

**Figure 1.**
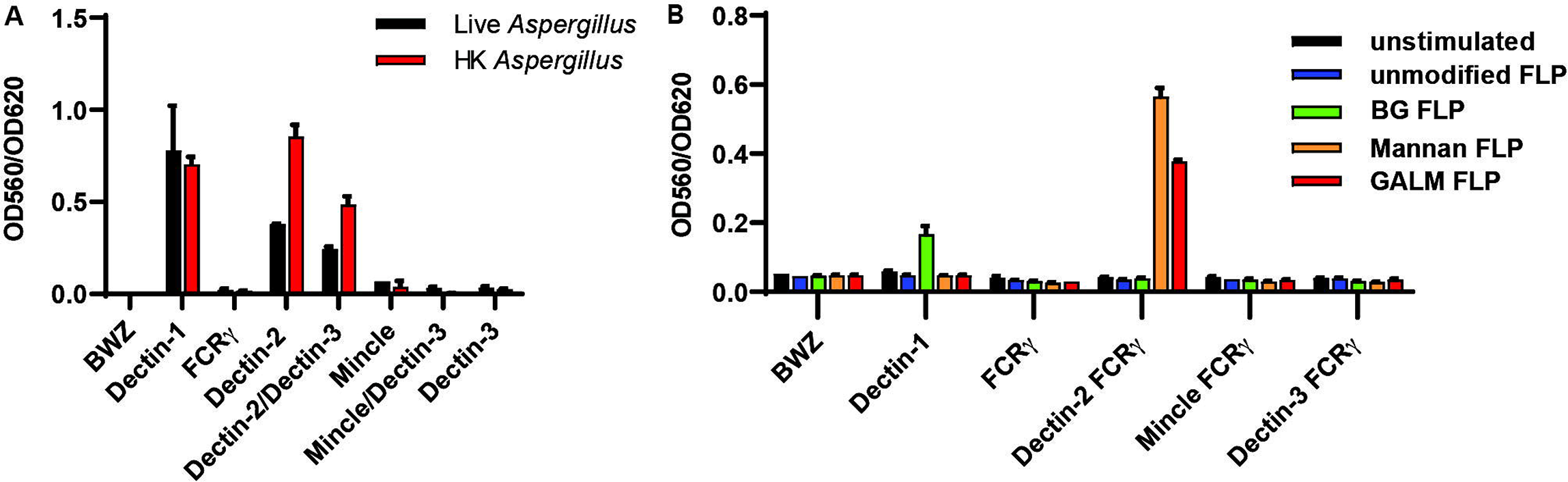
*A. fumigatus* and galactomannan activate Dectin-2 expressing CLR reporter cells. Reporter cells were stimulated for 18 hours using (A) either live or heat killed germlings of *A. fumigatus* Af293 at MOI 20:1 or (B) unmodified FLPs, β-1,3 glucan (BG) FLPs, *S. cerevisiae* mannan FLPs, and *A. fumigatus* galactomannan (GALM) FLPs at an effector to target ratio of 30:1. LacZ activity was measured in total cell lysates using CPRG as a substrate. Error bars represent the SEM of duplicate wells, and each experiment was repeated three times.

To dissect individual carbohydrate contributions to CLR reporter cell activation, we employed a technique developed in our laboratory for creating FLPs that consist of polystyrene beads coated with a discrete, purified fungal carbohydrates (34). We have previously generated FLPs made from purified β-1,3 glucan and demonstrated that carbohydrate-coated FLPs mimic cell-associated or particulate carbohydrates. These β-1,3 glucan FLPs can activate Dectin-1 signaling and elicit the formation of a phagocytic synapse (20, 34, 35). Generation of galactomannan FLPs was performed using methods previously described for the creation of β-1,3 glucan and mannan FLPs (34) (Supplemental Methods). All generated FLPs underwent rigorous quality control using flow cytometry to demonstrate the stable attachment of carbohydrates to the FLP surface (Figure S1). As an added control, multiple independent purifications of the galactomannan carbohydrate were used for FLP creation for all experiments and yielded similar results. We stimulated the reporter cell library with unmodified beads, β-1,3 glucan FLP, mannan FLP, and galactomannan FLP. Activation of CLR signaling resulted in expression of a LacZ reporter and was quantified using a CPRG assay to measure β-galactosidase activity. As expected, β-1,3 glucan FLPs activated Dectin-1 expressing reporter cells and *S. cerevisiae* mannan FLPs stimulated Dectin-2 expressing reporter cells. The galactomannan FLPs also stimulated Dectin-2 (Figure 1B) and Dectin-2/Dectin-3 co-expressing cells (data not shown), suggesting that not only did galactomannan bind to Dectin-2, but also triggered signaling by the CLR. None of our FLPs stimulated Mincle or Dectin-3 signaling alone, however these cells were activated by incubation with trehalose-6,6-dibehenate (TDB) (Figure S2) demonstrating that the receptors are functional, and that absence of observed stimulation is due to lack of CLR activation by the galactomannan FLPs. These data suggested that Dectin-2, but not Dectin-3 or Mincle, is a receptor for *Aspergillus*-derived galactomannan.

### Soluble Dectin-2 binds to galactomannan FLPs

After identifying Dectin-2 as a receptor for galactomannan using the CLR reporter cell screen, we next assessed whether Dectin-2 binds directly to galactomannan FLP. To evaluate this, we used soluble murine Dectin-2-human IgG1 Fc fusion protein and a control human IgG1 Fc protein (lacking Dectin-2). Flow cytometry using an anti-human IgG1 Fc antibody and an AF488 labeled secondary antibody was used to detect adherence of Dectin-2-Fc to the surface of FLP (Figure 2A). Dectin-2-Fc was specifically bound to both *Aspergillus* galactomannan FLPs and *Saccharomyces* mannan FLPs but did not bind to either unmodified or β-1,3 glucan FLPs. As expected, human IgG1 Fc protein did not bind to any FLP. As an additional control to demonstrate that binding to the galactomannan FLPs was Dectin-2 specific, we incubated the Dectin-2-Fc protein with anti-Dectin-2 neutralizing antibody or isotype control for 30 minutes prior to incubation with galactomannan FLPs. The Dectin-2 neutralizing antibody, but not the isotype control, potently blocked binding of Dectin-2-Fc to galactomannan FLP demonstrating a specific interaction (Figure 2B).

**Figure 2.**
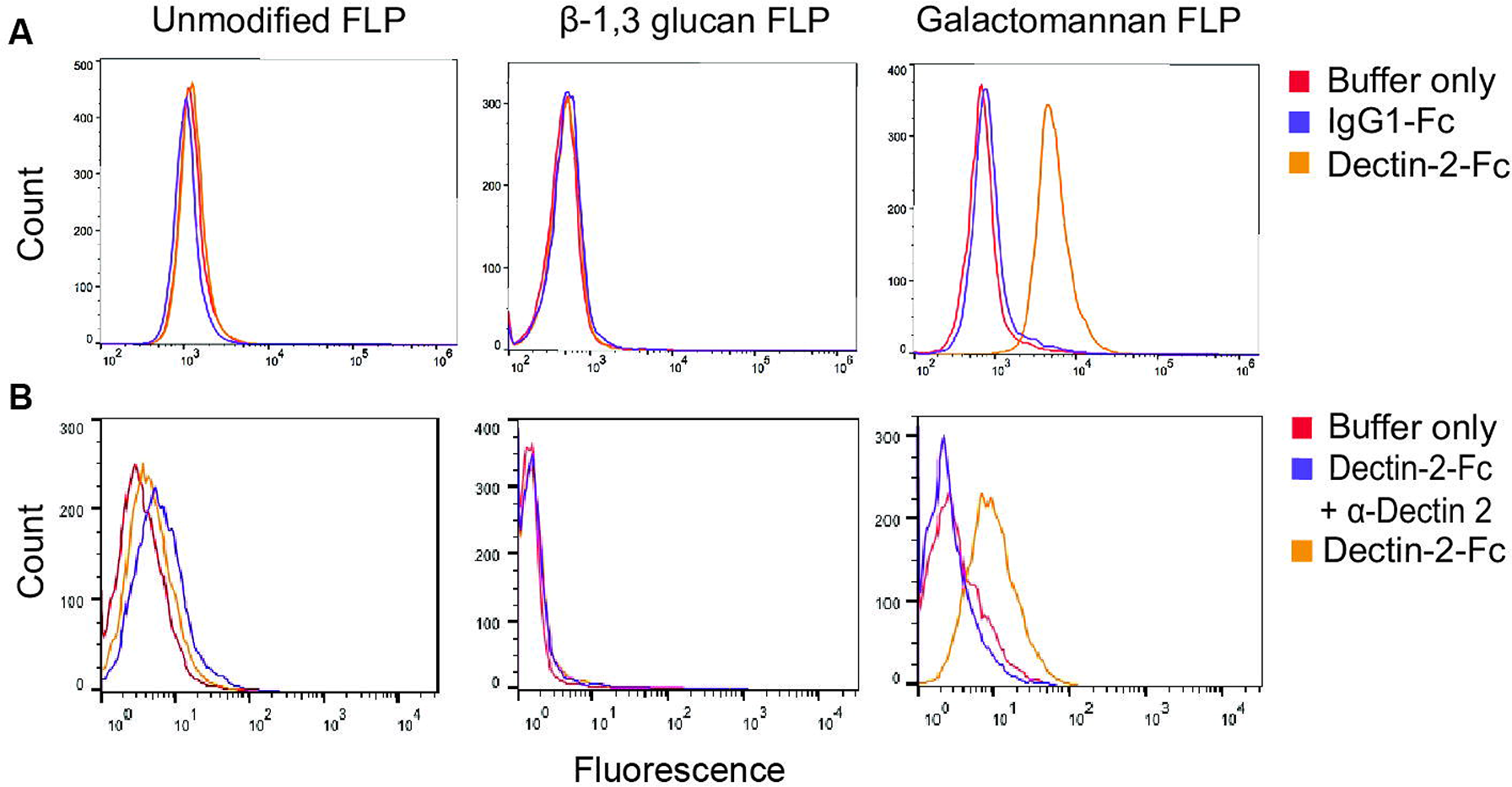
Recombinant Dectin-2 Fc protein binds galactomannan FLPs. (A) Unmodified FLPs, β-1,3 glucan FLPs, and galactomannan FLPs were incubated with soluble recombinant murine Dectin-2-human IgG1 Fc protein conjugate, control IgG1-Fc protein, or in buffer only, incubated with anti-human IgG1 conjugated to Alexa Fluor 488 and analyzed by flow cytometry. Dectin-2-Fc bound to galactomannan FLPs, but not to either unmodified or β-1,3 glucan FLPs. (B) FLPs were incubated for 2 hours in binding buffer alone or containing Dectin-2-Fc protein or Dectin-2-Fc pre-incubated for 30 minutes with a Dectin-2 neutralizing antibody. Dectin-2 neutralizing antibody disrupted the association of Dectin-2-Fc with galactomannan FLPs.

### Galactomannan FLPs induce Dectin-2 dependent Syk activation

The CLR reporter assay and binding studies demonstrated that Dectin-2 is capable of binding *Aspergillus* galactomannan. Next, we investigated whether this binding results in downstream signaling. Activation of Dectin-2 through ligand binding triggers phosphorylation of the ITAM motif of FcRγ and subsequent signaling through Syk. FcRγ is also required for localization of Dectin-2 to the cell surface (36). Macrophages express Dectin-2, however immortalized C57BL/6 murine macrophages have low levels of Dectin-2 surface expression at baseline (data not shown). Therefore, we transduced macrophages with lentivirus containing the murine Dectin-2 receptor (Supplemental Methods). As macrophages produce abundant FCRγ, we did not observe any difference in Dectin-2 surface expression between macrophages transduced with only Dectin-2 or those transduced with both Dectin-2 and FCRγ. To determine if galactomannan is sufficient to trigger Dectin-2 dependent Syk activation, we stimulated macrophages with galactomannan FLPs for 1 hour as well as unmodified FLPs, β-1,3 glucan FLPs, mannan FLPs, and *C. albicans.* We used *C. albicans* as a positive control in these experiments since it is a strong activator of Syk phosphorylation in wild-type macrophages. Lysates from stimulated cells were immunoblotted for phosphorylated and total Syk. As expected, we observed phosphorylation of Syk when wildtype macrophages were stimulated by *Candida albicans* and β-1,3 glucan FLPs (positive controls for Syk stimulation) (Figure 3A). There was increased phosphorylation of Syk in response to both galactomannan and mannan in Dectin-2 expressing macrophages demonstrating that *Aspergillus* galactomannan is sufficient to drive Dectin-2 dependent Syk phosphorylation (Figure 3A).

**Figure 3.**
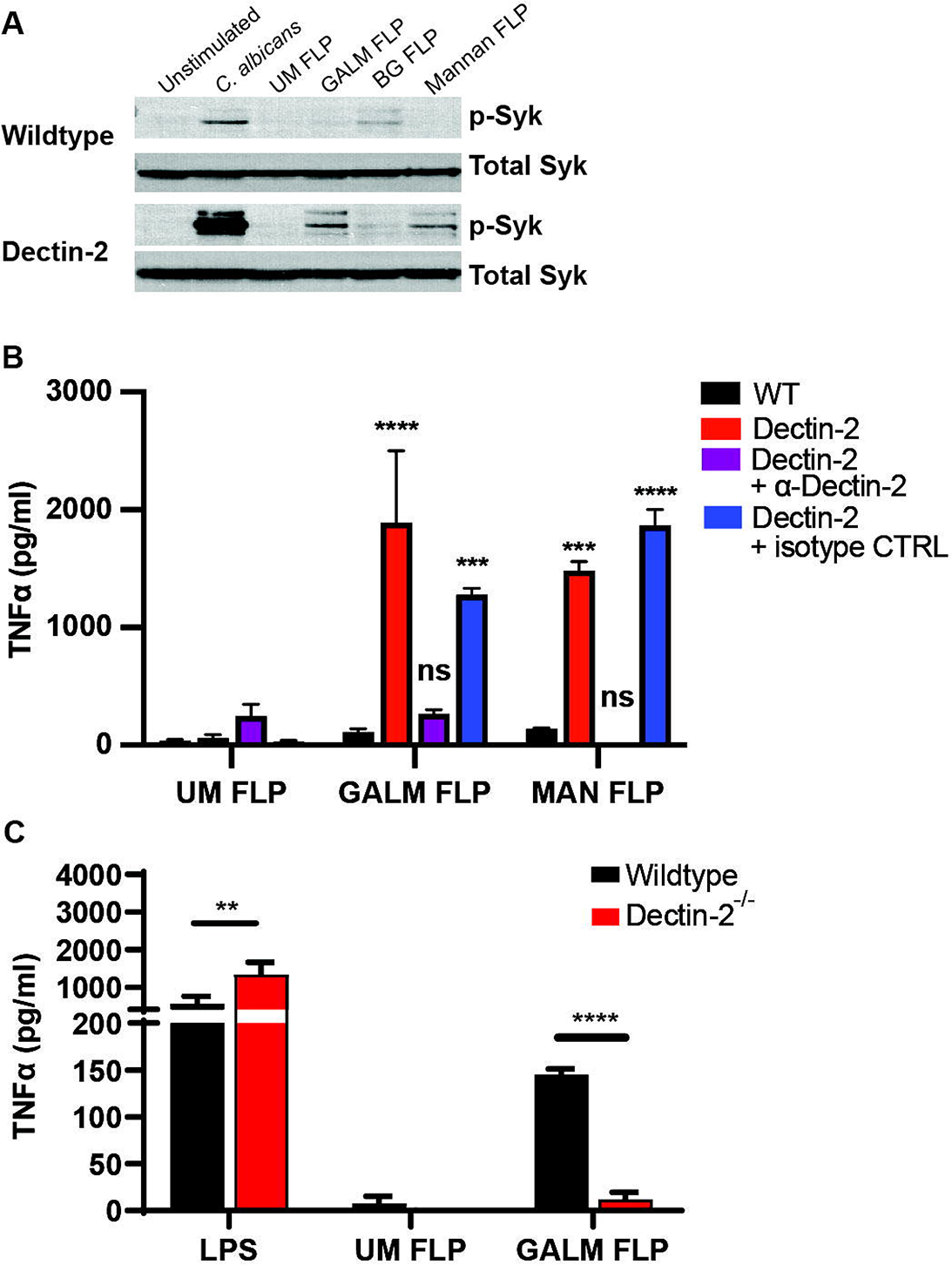
Galactomannan FLPs activate Dectin-2 dependent signaling in macrophages. (A) Immunoblot of immortalized wildtype C57BL/6 macrophages or C57BL/6 macrophages expressing Dectin-2 stimulated with live *C. albicans* (SC5314), unmodified FLPs (UM FLP), galactomannan FLPs (GALM FLP), β-1,3 glucan FLPs (BG FLP), or mannan FLPs at MOI 10:1 for 1 hour at 37°C and probed for both total Syk and phosphorylated (active) Syk. Overexpression of Dectin-2 increased phosphorylation of Syk in response to *C. albicans*, galactomannan and mannan FLPs, but not to unmodified or β-1,3 glucan FLPs. (B) Immortalized wildtype C57BL/6 macrophages or C57BL/6 macrophages expressing Dectin-2 were stimulated overnight with unmodified (UM), galactomannan (GALM), or mannan (MAN) FLPs at a target to effector ratio of 10:1. Prior to FLPs stimulation, macrophages were pre-treated for 30 minutes with either a Dectin-2 neutralizing antibody or an isotype control antibody as indicated. Supernatants were analyzed for TNFα using ELISA. Both galactomannan and mannan FLPs induced robust TNFα production by macrophages expressing Dectin-2, which was blocked by Dectin-2 neutralizing antibody, but not the isotype control. The significance indicated is in relation to the wild-type cell stimulated with the same FLPs. (C) BMDMs from wild-type C57BL/6 and Dectin-2^-/-^mice were stimulated with LPS (500 ng/mL), and unmodified (UM) or galactomannan (GALM) FLPs at an effector to target ration of 20:1 for 18 hours. Supernatants were analyzed for TNFα using ELISA. Dectin-2^-/-^ BMDM produced less TNFα when stimulated with galactomannan FLP compared with wildtype BMDM. Error bars represent standard error of three biological replicates, statistics were calculated with two-way ANOVA using PRISM 9 software (not significant, ns, *** p<0.001, **** p<0.0001).

### Dectin-2 mediates TNFα production by murine macrophages in response to both A. fumigatus *and galactomannan FLPs*

Having demonstrated that galactomannan is sufficient to activate Dectin-2 dependent Syk signaling, we next interrogated the downstream effects of this activation. Specifically, we sought to determine if galactomannan stimulation of Dectin-2 is both sufficient and necessary for cytokine production. To determine if galactomannan stimulation of Dectin-2 is sufficient to trigger cytokine production, Dectin-2 expressing murine macrophages were stimulated overnight with unmodified FLPs, β-1,3 glucan FLPs, *A. fumigatus* galactomannan FLPs, or *S. cerevisiae* mannan FLPs (positive control) and TNFα production was measured by ELISA. Galactomannan and mannan FLPs enhanced TNFα production in Dectin-2 expressing macrophages (Figure 3A). To demonstrate that this effect was specific to Dectin-2, we pre-incubated the macrophages with a Dectin-2 neutralizing antibody or an isotype control for 30 minutes prior to stimulation with FLPs. TNFα production was blocked by the Dectin-2 specific antibody, but not by the isotype control, demonstrating a specific role for Dectin-2 in mediating the TNFα production in response to galactomannan (Figure 3B).

To examine if Dectin-2 is required for primary BMDMs to respond to galactomannan FLPs, primary BMDM from wildtype or Dectin-2 deficient mice were stimulated by galactomannan FLPs, unmodified FLPs (negative control), or LPS (positive control). There was near complete reduction of TNFα production in response to galactomannan FLPs (Figure 3C), indicating that galactomannan triggers the secretion of pro-inflammatory cytokines in BMDMs in a Dectin-2 dependent manner.

### Deletion of Dectin-2 results in increased recruitment of immune cells, but does not alter fungal burden

We sought to determine if the immune cell influx in response to infection was altered in the absence of Dectin-2. Since treatment with steroids affects immune cell function, we chose to perform this experiment in an immunocompetent, rather than a steroid-induced immunosuppression mouse model. Immunocompetent mice were administered either *A. fumigatus* strain CEA10 or PBS via oropharyngeal aspiration. After 48 hours of infection lungs were harvested, digested, and the resulting single cell suspension was enriched for leukocytes using a Percoll gradient. A multi-channel flow cytometry experiment was performed to allow simultaneous assessment of multiple cell lines including neutrophils, eosinophils, T cells, B cells, dendritic cells (including cDC1, cDC2, monocyte derived DCs, and plasmacytoid dendritic cells [pDCs]), NK cells, monocytes, and alveolar macrophages. All infected mice demonstrated a marked increase in CD45^+^ cellular infiltrates compared to the PBS control mice. We infected mice with 1 x 10^7^, 2 x 10^7^ and 4 x 10^7^ conidia of CEA10 *A. fumigatus* and noted a dose-dependent increase in immune cell infiltrate. At doses higher than 4 x 10^7^ most mice succumbed within 24 – 48 hours, presumably due to an exuberant inflammatory response. Infected Dectin-2^-/-^ mice recruited more CD45^+^ cells to the lungs than did infected wild-type mice. Additionally, Dectin-2^-/-^ infected mouse lungs had a significant increase in innate cells (Figure 4B, Supplemental Figure S3), as well as neutrophils compared to wild-type (Figure 4C). Notably, the proportion of the cellular infiltrate comprised of innate cells (Figure 4D) and neutrophils (Figure 4E) was not significantly different between infected wild-type and Dectin-2^-/-^ mice suggesting that there is increased immune cell recruitment to the lungs during infection in Dectin-2^-/-^ mice, but no change in the overall distribution of innate cell and neutrophil populations. Our results suggest that Dectin-2 is involved in the coordination of inflammatory responses in this airway model of infection.

**Figure 4.**
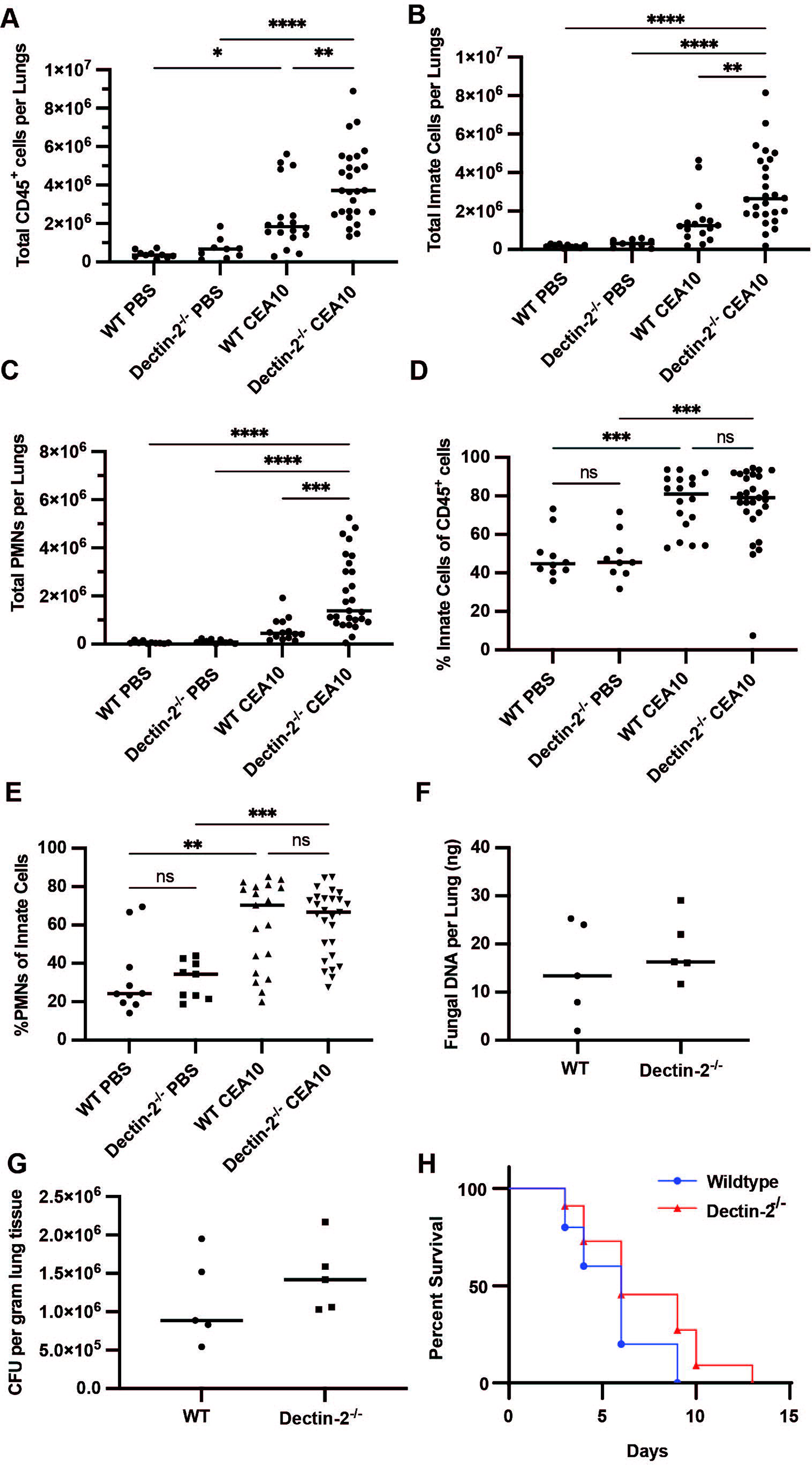
Deficiency of Dectin-2 leads to increased immune cell recruitment, but is dispensable for survival. (A - E) Immunocompetent wild-type and Dectin-2^-/-^ mice were infected via oropharyngeal inhalation of 4 x 10^7^ conidia of *A. fumigatus* CEA10. Lung tissue was harvested 48 hours post-infection and digested to obtain a single cell suspension that was then analyzed by flow cytometry. Dectin-2 infected mice had (A) increased infiltration of live CD45+ cells compared with wild-type. There was a significant increase in the total number of (B) innate cells (7-AAD^lo^, CD45^+^, CD90.2^-^, CD19^-^) and (C) neutrophils (7-AAD^lo^, CD45^+^, CD90.2^-^, CD19^-^, SiglecF^-^, Ly6G^+^) in Dectin-2^-/-^ mice compared with wild-type. However, there was no difference in the percentage of cells that were innate cells or (E) neutrophils. The data shown is a composite from two independent experiments, consisting of at least 5 PBS control mice and 10 CEA10 infected mice per mouse strain. Each point represents an individual animal and horizontal lines represent the median. One-way ANOVA was performed using PRISM9 software to determine statistical significance (not significant, ns, * p<0.05, ** p<0.01, *** p<0.001, **** p<0.0001). (F,G) Immunocompetent wild-type and Dectin-2^-/-^ mice were infected with 4 x 10^7^ CEA10. After 48 hours lungs were extracted, and fungal burden was quantified using (F) qPCR or (G) CFU. There was no difference in fungal burden between wild-type and Dectin-2^-/-^ mice. Each point represents an individual animal and horizontal lines represent the median fungal burden in the lungs. Data shown are from a single replicated consisting of 5 infected and 1 PBS control per mouse strain. Each experiment was repeated at least twice, and concordant results were obtained in each experiment. (H) Wild-type C57BL/6 and Dectin-2^-/-^ mice were immunosuppressed with corticosteroids and infected intranasally with 2.5 x 10^4^ conidia of *A. fumigatus* strain CEA10. There was no significant difference in survival between wildtype and Dectin-2^-/-^ mice as determined based upon Kaplan-Meyer and Cox Analysis using PRISM 9 software.

To determine whether the increased influx of CD45^+^ cells could be the result of difference in fungal burden between wild-type and Dectin-2 deficient mice, we quantified fungal burden in the lungs of mice infected with 4 x 10^7^ conidia at 48 hours post-infection using qPCR to detect and quantify *Aspergillus* DNA(37). *Aspergillus fumigatus* DNA was detected in all samples from infected mice, but not in PBS controls. There was no difference in *Aspergillus* DNA between wild-type and Dectin-2 deficient mice (Figure 4F). CFU assays were also performed as a confirmatory test to ensure that there was no difference in live *Aspergillus* fungal burden, since fungal DNA testing cannot distinguish between live versus dead organisms. Lungs from both wild-type and Dectin-2 deficient mice still contained live *Aspergillus fumigatus* at 48 hours, and there was no difference in CFU between strains (Figure 4G). Taken together these results suggest that the enhanced immune cell infiltration in Dectin-2 deficient mice is not due to a difference in fungal burden.

### Dectin-2 is dispensable for survival in a murine model of infection

Our data suggest that Dectin-2 is a receptor for *A. fumigatus* galactomannan, and that recognition of galactomannan by Dectin-2 results in Syk phosphorylation, cytokine production and increased inflammatory cell recruitment in Dectin-2^-/-^ mice infected with *A. fumigatus*. To determine if this pathway is required for survival during *Aspergillus* infection, we intranasally infected corticosteroid immunosuppressed Dectin-2^-/-^ and wild-type C57BL/6 mice with *A. fumigatus* (CEA10 strain) and monitored the mice for signs of infection. There was no difference in survival between wild-type and Dectin-2 deficient mice, suggesting that in this model, Dectin-2 is dispensable for survival (Figure 4H).

## DISCUSSION

Galactomannan is a clinically important polysaccharide released into patients’ blood and bronchial lavage fluid and used to diagnose invasive aspergillosis infections. However, we do not understand the immunological implications of the wide-scale release of this carbohydrate into patient tissue, or the role that it plays in mediating direct immunological interactions between *A. fumigatus* and innate immune cells at the site of infection. Here, we utilized FLPs coated with purified *A. fumigatus* galactomannan to identify Dectin-2 as a receptor for galactomannan. Our results demonstrate that cell-associated galactomannan binds Dectin-2 inducing phosphorylation of Syk and TNFα production.

Although BMDMs from Dectin-2 deficient mice had decreased cytokine production in response to galactomannan FLPs, there was no difference in overall survival of immunosuppressed Dectin-2 deficient mice compared with wild-type mice when challenged intranasally with *A. fumigatus*. There is significant redundancy in CLRs signaling, therefore it is possible that the lack of Dectin-2 is compensated for through Dectin-1 or other CLR-ligand interactions in the context of the whole organism. However, publications have suggested a critical role for Dectin-2 in the immune response to *A. fumigatus*(*38, 39*). A recent case report identified a patient without previously recognized immunodeficiency that developed invasive aspergillosis and was found to have a mutation in Dectin-2 that led to decreased responsiveness to *Aspergillus* (38). Previous work has also demonstrated a role for Dectin-2 in mediating plasmacytoid dendritic cell (pDC) responses to *A. fumigatus* (39). These observations, coupled with this study, suggest that while Dectin-2 may not be required for survival in a murine model, it may contribute to human disease through modulating interactions with discrete immune cell populations.

Interestingly, immunocompetent mice that lacked Dectin-2 have increased recruitment of immune cells into the lung tissue upon infection with *Aspergillus*, which is in sharp contrast to galactosaminogalactan (GAG) from *A. fumigatus* which led to a reduction of neutrophil infiltration (40). The difference in inflammatory cell infiltrates was not due to difference in the fungal burden within the lungs, suggesting that it is a direct result of Dectin-2 deficiency. This result is interesting, as our data also demonstrated that galactomannan-induced Dectin-2 signaling in macrophages leads to pro-inflammatory TNFα release. We anticipated that the absence of Dectin-2 would result in decreased pulmonary inflammation, but surprisingly observed increased immune cell recruitment. Our results suggest that Dectin-2 may normally temper inflammatory cell recruitment to the lungs, or that Dectin-2 signaling normally leads to increased inflammatory cell turnover. There are multiple potential hypotheses to explain these findings which will provide fertile ground for future studies. In the absence of Dectin-2, expression of Dectin-1 and/or other C-type Lectin Receptors may be upregulated and thus these receptors may play a more substantial role in driving the immune response. Additionally, Dectin-2 is expressed not just by macrophages, but also by many immune cells and other lung resident cells, including epithelial cells. Thus, a cell type other than macrophages, may be responsible for mediating the difference in immune cell influx that we observed. For instance, studies by Loures, *et al* (39) demonstrated that plasmacytoid dendritic cell production of type I IFNs in response to *Aspergillus fumigatus* is dependent upon Dectin-2 mediated recognition of *Aspergillus fumigatus*. While we did not observe a difference in pDC recruitment to the lungs in wild-type compared to Dectin-2 deficient mice, these cells are likely impaired in their ability to produce type I IFNs during infection. Interestingly, the increased immune cell influx observed in Dectin-2 deficient mice did not affect fungal burden suggesting, that the increased immune cells infiltrate did not result in more rapid clearance of infection. Whether the increased inflammatory influx seen in Dectin-2 deficient mice could have detrimental effects on the host, such as increasing lung damage during infection is unknown.

Our current work addresses the mechanism by which cell-wall associated galactomannan is recognized by Dectin-2. Galactomannan and β-1,3 glucan are both fungal cell wall carbohydrates that exists in both soluble and cell-wall associated states. β-1,3 glucan is known to interact with the immune system differently depending upon its form. Whether galactomannan-induced immune activation is triggered only at sites of infection or more systemically and how this could impact the outcome of infection remain critically important questions. Whether circulating soluble galactomannan can stimulate recognition by Dectin-2 will be the subject of future studies. Dectin-2 has also been implicated in allergic disease (41–44), suggesting that galactomannan from *Aspergillus* could play a role in stimulating allergic manifestations to this pathogen. Overall, our data demonstrate that galactomannan is recognized by Dectin-2 and lays the exciting groundwork for dissecting the role of galactomannan in an important clinical disease.

## MATERIALS AND METHODS

### Animals

Mice were maintained in specific-pathogen-free barrier facilities at Massachusetts General Hospital (MGH; Boston, MA) according to Institutional Animal Care and Use Committee (IACUC) Guidelines. Wildtype C57BL/6 mice were obtained from Jackson Laboratories (Bar Harbor, ME) and Dectin-2^-/-^ mice (45) were from the University of Wisconsin-Madison. All mice used for bone marrow harvests and *in vivo* experiments were 8 – 20 weeks of age and were co-housed prior to experiments. Mice were both age and sex-matched for each experiment and an equal distribution of male and female mice used.

### Cell lines and culture

Immortalized murine C57BL/6 bone marrow-derived macrophages (BMDM) were a gift from Doug Golenbock (University of Massachusetts Medical School, Worcester, MA). Macrophages were cultured in complete RPMI media (cRMPI; RPMI 1650, L-glutamine, 10% heat-inactivated fetal bovine serum (FBS; Gibco, Thermo Fisher Scientific, Rockford, IL), 1% penicillin/streptomycin, 1% HEPES buffer, and 50 μM of β-mercaptoethanol). Puromycin and blastocidin were added to a final concentration of 5 μg/mL as needed for the selection of transduced cells. Primary BMDMs from C57BL/6 and Dectin-2^-/-^ mice were harvested as previously described (35). After harvest, primary BMDMs were grown for 7 days in complete RPMI media containing 20 ng/mL of recombinant MCSF (Peprotech, Rocky Hill, NJ) prior to use in assays. HEK293T cells were purchased from the ATCC (American Type Cell Collection, Manassas, VA) and were grown in complete DMEM (DMEM, L-glutamine, 10% heat-inactivated FBS, 1% penicillin/streptomycin, 1% HEPES buffer). All cell lines were grown at 37^0^C in the presence of 5% CO_2_.

### Fungal culture

*A. fumigatus* strains Af293 and CEA10 were grown at 37℃ for 5 days on Sabouraud’s Dextrose Agar (SBD; Difco, Fisher Scientific, Rockford, IL) or for 3 days on glucose minimal media slants (GMM), respectively. Slants were seeded from frozen stocks of conidia maintained at −80℃. To harvest conidia, a sterile solution of deionized water (Milli-Q, Millipore Sigma, Burlington, MA) containing 0.01% Tween 20 was added to each slant and spores were liberated using gentle surface agitation with a sterile swab. The spore solution was passed through a 40 μm cell strainer to separate hyphal debris. Spores were washed three times with sterile PBS and counted on a LUNA^TM^ automated cell counter (Logos Biosystems, Annandale, VA) and diluted appropriately. To generate *A. fumigatus* germlings, conidia were incubated in cRPMI media at 37℃ for 4-6 hours and visually inspected for the presence of emerging germ tubes. Germlings were then washed, counted, and resuspended in PBS at the desired inoculum. For heat-killing, *A. fumigatus* germlings were incubated at 95℃ for 30 minutes.

### Purification of galactomannan

The galactomannan was isolated as lipogalactomannan from the cellular membrane of *A. fumigatus* mycelium that had been grown in SBD medium using previously described methods (27). The lipid moiety was removed by nitrous deamination and the polysaccharide moiety was purified by gel filtration chromatography on Superdex 75 column (10/300 GL, GE-Healthcare) (27). Three independent preparations of galactomannan were used to create FLP and multiple batches were used per experiment to confirm that results were not batch dependent.

### FLP binding assays

FLPs were blocked overnight in PBS containing 2% BSA. FLPs were washed and resuspended in 100 mL of binding buffer (20 mM Tris-HCL, pH 7.4, 150 mM NaCl, 10 mM CaCl_2,_ 0.05% Tween-20). Dectin-2(murine)-IgG1 Fc (human) fusion protein (ENZO Life Sciences, Inc, Farmingdale, NY) or IgG1 Fc (human) protein (Thermo Fisher Scientific, Rockford, IL) was added to a final concentration of 10 mg/mL and samples were incubated with gentle mixing for 2 hours at room temperature. Following washing, samples were resuspended in binding buffer containing a 1:100 dilution of mouse anti-human-IgG1-Fc conjugated to Alexa Fluor 488 (Thermo Fisher Scientific, Rockford, IL; A-10631) and incubated for 30 minutes at room temperature in the dark. FLPs were washed with binding buffer, resuspended in FACS buffer (PBS containing 2% BSA), examined on a BD FACSCalibur^TM^ or BD FACSCelesta^TM-^ Flow Cytometer (BD, Franklin Lakes, NJ), and data analyzed using FlowJo 10. For neutralizing antibody experiments, mouse anti-Dectin-2 antibody (R&D Systems, MAB1525) was preincubated with the Dectin-2-Fc fusion protein at a concentration of 50 mg/ml for 30 minutes prior to adding the mixture to FLPs.

### In vivo Aspergillus survival model

Wild-type C57BL/6 mice or Dectin-2^-/-^ mice were immunosuppressed with 40 mg/kg subcutaneous triamcinolone acetonide (Kenalog^®^-10, Bristol-Myers Squibb, New York, NY) starting the day prior to infection and every 7 days thereafter. The mice were infected with 2.4 x10^4^ CEA10 conidia intranasally under isoflurane anesthesia. For each experiment 10 mice were infected with CEA10 and 3 mice were treated with PBS only per genotype. Mice were monitored twice daily for the first 5 days of the experiment and then daily thereafter. Mice were assessed based upon a 12-point scale to determine when an individual animal should be euthanized due to illness and removed from the study. This was developed in conjunction with the MGH IACUC and includes the following symptoms and points assigned to each in parentheses: hunched posture (3), ruffled and/or matted fur (3), shivering (3), abnormal breathing (increased respiratory rate) (12), 75% reduction in activity compared to controls (3), inactivity leading to inability to acquire food or water (12), and barrel rolling (12). If any animal reaches 12 points, they are humanely euthanized and counted as a death for the purposes of survival analysis. All animal experiments were approved under the MGH IACUC protocol #2008N000078.

### Western blot

Cells were placed on ice and lysed with 1% NP40 lysis buffer containing protease and phosphatase inhibitors. Proteins were denatured using NuPAGE LDS Sample Buffer (Thermo Fisher Scientific, Rockford, IL). The proteins were resolved by SDS-PAGE on 4-12% NuPAGE gels using NuPAGE MOPS buffer (NuPage gels, Thermo Fisher Scientific, Rockford, IL), and transferred to methanol-activated PVDF membrane (Perkin Elmer, Waltham, MA) using transfer buffer (0.025M Tris, 0.192 M glycine, 20% methanol) and electrophoretic transfer at 100V for 1 hour. For detection of proteins, PVDF membranes were blocked for 1 hour at room temperature in 5% BSA in PBS-0.01% Tween 20 (PBST). Blots were incubated with anti-phospho spleen tyrosine kinase (Syk) antibody (Cell Signaling Technologies, Danvers, MA; 2710S) or anti-total Syk antibody (Cell Signaling Technologies, 13198S), in 5% BSA in PBST overnight at 4℃. The membranes were washed and incubated with secondary swine anti-rabbit HRP conjugated antibody (Agilent DAKO, Santa Clara, CA, P0399) in 1% milk in PBST for 1 hour at room temperature. Membranes were washed and then visualized using Western Lightning Plus ECL chemiluminescent substrate (Perkin Elmer, Waltham, MA) on Kodak BioMax XAR film (MilliporeSigma, Burlington, MA). Films were then scanned and processed using Adobe Photoshop 2021. Any contrast adjustments were applied evenly to the entire image and adheres to standards set forth by the scientific community (46).

## ACKNOWLEDGEMENTS

We thank all the members of the Vyas and Mansour lab at MGH and Dr. Vishukumar Aimanianda, Institut Pasteur, for useful discussions and reading of the manuscript.

## AUTHOR CONTRIBUTIONS

Conceptualization – Jennifer Reedy and Jatin Vyas. Investigation – Jennifer Reedy, Arianne Crossen, Paige Negoro, Marcel Wuethrich, Kyle Timmer, and Hannah Brown. Writing – Jennifer Reedy and Jatin Vyas, Manuscript review and revision – Arianne Crossen, Paige Negoro, Kyle Timmer, Hannah Brown, Diego Vargas Blanco, Rebecca Ward, Thierry Fontaine, Marcel Wüthrich, Michael Mansour, Jatin Vyas. Resources - Thierry Fontaine, Marcel Wuethrich, Michael K. Mansour, and Jatin Vyas.

## SUPPLEMENTAL MATERIALS

**Supplemental Methods:** Generation and quality control of FLP, Lentiviral transduction of cells with Dectin-2 and FcRγ, CLR ligand reporter cell assay, ELISA, Immunophenotyping of pulmonary infiltrates, quantification of fungal burden in lungs, Colony forming unit (CFU)

**Figure S1:** Generation and validation of galactomannan FLP

**Figure S2:** CLR reporter cell activation using CLR control agonists

**Figure S3:** Flow cytometry gating strategy for lung immunophenotyping to identify neutrophils.

