## supplemental Methods, Figures S1,S2,S3 for "The C-type Lectin Receptor Dectin-2 is a receptor for *Aspergillus fumigatus* galactomannan"

### *Generation and quality control of FLP*

Carbohydrate coated-FLP consisting of purified $\beta$-1,3 glucan (Wellmune, Biothera, *Saccharomyces cerevisiae*, Eagan, MN) and mannan (*S. cerevisiae*, Sigma-Aldrich, M704-1G) were creating using previously published methods (1). Using purified *Aspergillus fumigatus* galactomannan, we created both adsorbed FLP in which the carbohydrate is attached to the bead through electrostatic interactions and covalently-attached FLP using 1,1'-carbonyldiimidazole (CDI)-chemistry as previously published (1). Briefly, 300 mL of amine-coated polystyrene beads (Polysciences, Warrington, PA) were washed three times in sterile anhydrous dimethylsulfoxide (DMSO, MilliporeSigma, Burlington, MA) using 0.45 $\mu$m PTFE centrifugal filters (MilliporeSigma, Burlington, MA) to remove any aqueous solution. The beads were then resuspended in either DMSO (adsorbed beads) or DMSO containing 0.5 M 1,1'-carbonyldiimidazole (CDI) and incubated at room temperature for 1 hour with shaking. Beads were then rinsed with DMSO and resuspended in 1mg/mL purified *A. fumigatus* galactomannan in DMSO. Unmodified beads were resuspended just in DMSO without additional carbohydrates. The mixture was incubated for 1 hour at room temperature with shaking and centrifuged to remove the DMSO and excess carbohydrates. Beads were then washed, and resuspended in PBS, counted on a Luna™ automated cell counter, and stored at 4°C.

To confirm the association of carbohydrates with the FLPs and ensure rigor, each newly created batch of FLPs was analyzed by flow cytometry. Briefly, 3 mL of beads were blocked overnight in PBS containing 2% BSA at 4℃. To analyze mannan binding to the surface, 10 mL of 20 mg/mL stock fluorescein labeled Concanavalin A (Con A; Invitrogen, Thermo Fisher Scientific, Carlsbad, CA; C827) was added to each sample and incubated for 1 hour at room temperature while shaking. Samples were washed and resuspended in FACS buffer (PBS containing 2% BSA) and then analyzed on a BD FACSCalibur™ or BD FACSCelesta™ Flow Cytometer (BD, Franklin Lakes, NJ). For analysis of β-1,3 glucan, 20 ng of primary monoclonal anti-β-1,3 glucan antibody (Biosupplies Australia Pty LTD, Victoria, Australia; 400-2) was added to each sample and incubated for 1 hour at room temperature. FLPs were washed and then incubated for 30 minutes with a 1:100 dilution of secondary rabbit anti-mouse IgG conjugated with Alexa Fluor 488 (Invitrogen, Carlsbad, CA; A11059). FLPs were then washed, resuspended in FACS buffer, and analyzed by flow cytometry. For galactomannan-coated beads, FLPs were centrifuged, and the blocking buffer was removed. FLPs were resuspended in 50 ml of component R6 from the Platelia *Aspergillus* Ag kit (Bio-Rad, Hercules, CA) containing rat anti-galactomannan IgM antibody and were incubated for 1 hour at room temperature. FLPs were then washed and resuspended in a 1:200 dilution of secondary goat anti-rat IgM conjugated with Alexa Fluor 488 (Invitrogen, Carlsbad, CA; A-21212) in FACs Buffer. FLPs were incubated for 30 minutes at room temperature and then washed and resuspended in FACS buffer.

To assess the stability of carbohydrate association, 10 mL of FLP were incubated in 800 mL of either 3% SDS, 1X NP40, 1% Triton X-100 for 1 hour at room temperature with constant shaking, or 3% SDS at $95^0$C for 1 hour. The FLPs were centrifuged, washed with detergent, and washed three times with FACS buffer. FLPs were then subjected to antibody staining to assess for carbohydrate association as detailed above.

For fluorescent imaging of carbohydrate coated FLPs, FLPs were fluorescently labeled in the same manner as for flow cytometry and then placed onto glass slides. FLPs were imaged using a Nikon Ti-E inverted microscope with a CSU-X1 confocal spinning-disk head (Yokogawa, Sugarland, TX) with a Coherent 4-Watt laser (Coherent, Santa Clara, CA) as the excitation source. Images were acquired through an EM-CCD camera (Hamamatsu, C9100-13, Bridgewater, NJ) and performed using MetaMorph® software (Molecular Devices, Downington, PA). Image data files were processed using Adobe Photoshop 2021 and assembled in Adobe Illustrator 2022 (Adobe Systems, San Jose, CA).

***Lentiviral transduction of cells with Dectin-2 and FcRy***

Murine Dectin-2 was amplified from plasmid pFB-dectin2-IRES-eGFP from Caetano Reis e Sousa using a 5' primer containing NotI digestion site with sequence TATTGCGGCCGCCATGGTGCAGGAAAGACAATCCCAAGG and a 3' primer containing a BglII restriction site with sequence TCGACGATAGATCTTCATAGGTAAATCTTCTTCATTTCACATATTGAATTG. Primers were synthesized by Integrated DNA technologies (IDT, Research Triangle Park, NC). The gamma chain protein (FcRγ) of the Fc-gamma receptor (FcγR) was PCR amplified from plasmid pSVL-mo FcεRI gamma (plasmid was a gift from Jean Kinet, Addgene plasmid #8372), using 5' primer containing a NotI digestion site with sequence TATTGCGGCCGCCATGATCTCAGCCGTGATCTTGTTCTTG and 3' primer containing a BamHI digestion site TCGACGATGGATCCCTACTGGGGTGGTTTTTCATGCTTC. The constructs were cloned into a pHAGEII lentiviral vector containing either puromycin or blasticidin selectable markers using NotI and BamHI digestion and sequence verified. Generation of lentivirus in HEK293T cells and transduction was performed as previously described (2). Cells were selected for uptake of the lentivirus vector using puromycin or blasticidin at 5 μg/mL. After selection, expression of the desired gene was confirmed using western blot. Surface localization of Dectin-2 was confirmed using flow cytometry using anti-murine Dectin-2-APC (R&D Systems, Minneapolis, MN, FAB1515A) or anti-human Dectin-2 polyclonal goat antibody (R&D, AF3114) with Donkey anti-Goat AF488 secondary antibody (Invitrogen, Thermo Fisher Scientific, Rockford, IL, A11055).

***CLR ligand reporter cell assay***

Reporter cells were used based on previously published protocols (3). B3Z and BWZ cells carrying an NFAT-lacZ reporter construct have been previously described (4, 5). B3Z cells expressing murine Dectin-2 or a wildtype murine FCRγ-chain as well as BWZ cells expressing murine Dectin-1CD3ζ (the extracellular domain of mouse Dectin-1 fused to CD3ζ intracellular tail) were constructed by Dr. Caetano Reis e Sousa (London Research Institute, London, U.K) (6). Mincle and Dectin-3 (MCL) expressing cells were created as previously described (3). Briefly, cells were seeded at $1 \times 10^5$ cells per well in 96-well plates followed by stimulation with FLPs or media alone. Cells were incubated for 18 hours in a tissue culture incubator at 37℃ with 5% $CO_2$. Cells were pelleted and the supernatant was removed. Cells were lysed with triton lysis buffer (1% Triton X-100, 9 mM $KH_2PO_4$, 90 mM $K_2HPO_4$) on ice for 30 minutes. Plates were centrifuged to pellet cellular debris and 40 mL of lysate was transferred to a new 96-well plate. Assay buffer containing chlorophenol red-β-D-galactopyranoside (CPRG) was added to each well and plates were incubated for 6 hours at 37℃ protected from light and then absorbance at 560 nm and 620 nm was measured using an i3x Spectrophotometer (Molecular Devices LLC, San Jose, CA).

***ELISA***

1 x$10^5$ murine macrophages (immortalized or BMDM) were plated in triplicate in tissue culture treated 48-well plates and stimulated overnight with FLP or fungal germlings at the indicated target-to-effector ratios. Supernatants were collected for cytokine analysis using ELISA (Duoset, R&D Systems, Minneapolis, MN) and read using an i3X Spectrophotometer (Molecular Devices, LLC, San Jose, CA). Results were analyzed using PRISM9 software (GraphPad Software, San Diego, CA).

***Immunophenotyping of pulmonary infiltrates***

Wildtype or Dectin-$2^{-/-}$ mice were infected with 4 x $10^7$ conidia of *A. fumigatus* strain CEA10 through oropharyngeal inhalation. Each experiment consisted of at least 10 wild-type and 10 Dectin-$2^{-/-}$ mice infected with CEA10 *Aspergillus* and 5 mock infection (PBS only) control per mouse line. Mice were euthanized at 48 hours post-infection and immunophenotyping was performed as previously described(7). Briefly, lungs were perfused with PBS through cardiac puncture to flush intravascular cells. Luns were excised, placed into Digestion Solution (38 mg Collagenase Type I and 100 U DNase I in 100 ml RPMI 1640), finely minced using scissors and incubated for 1 hour with shaking at 37℃. After 1 hour, 800 ml of 0.5 M EDTA was added to each sample to stop the collagenase activity. Lungs samples were passed through a 70 mM filter, pelleted, resuspended in a 40% Percoll-RPMI solution, and under-layered with a 67% Percoll-RPMI solution. Samples were centrifuged at 650 x g for 20 minutes and leukocytes were collected at the interface between layers and washed twice with FACS buffer (PBS with 2% FBS). Cells were counted and 1 – 2 x $10^6$ cells were stained for flow cytometry analysis. Fc block (anti-Mouse CD16/CD32, Thermo Fisher Scientific, Rockford, IL, 14-0161-85) was added to cells at 1:100 dilution and incubated at room temperature for 15 minutes. Cells were then labeled with a mixture of the following fluorophore-conjugated antibodies for 1 hour at 4℃ (all antibodies are from Biolegend, San Diego, CA unless otherwise noted): anti-CD45 BV605 (Biolegend, 103139), anti-CD90.2 BV786 (Biolegend, 105331), anti-CD19 PE-Dazzle (Biolegend, 115553), anti-CD4 BV510 (Biolegend, 100559), anti-CD8 APC FIRE (Biolegend, 100765), anti-Siglec-F APC (Biolegend, 155507), anti-Ly6G AF 488 (Biolegend, 127625), anti-Ly6C BV510 (Biolegend, 128033), anti-I-A/I-E AF700 (Biolegend, 107622), anti-NCR1 BV421 (Biolegend, 137611), anti-CD64 PE (Biolegend, 139303), anti-CD103 BV421 (Biolegend, 121421), anti-CD11b APC FIRE (Biolegend, 101261), anti-CD11c BV650 (Biolegend, 117339), anti-B220/CD45R PE (Biolegend, 103207), anti-TCRγδ APC (Biolegend, 118115), and anti-TCRβ BV650 (Biolegend, 109251). Cells were washed with FACS buffer, and 7-AAD (Stem Cell Technologies, 75001.1) was added for live-dead cell staining. Flow cytometry was performed using BD FACSCelesta™ and analysis was performed using FlowJo 10 software (BD, Ashland, OR).

**Quantification of Fungal Burden in Lungs**

Wildtype or Dectin-2$^{-/-}$ mice were infected with 4 x $10^7$ conidia of *A. fumigatus* strain CEA10 through oropharyngeal inhalation under isoflurane anesthesia**.** Each experiment consisted of 5 wild-type and 5 Dectin-2$^{-/-}$ mice infected with CEA10 *Aspergillus* and one mock infection (PBS only) control per mouse line. After 48 hours lungs were extracted, frozen at -80ºC, and lyophilized overnight. Freeze-dried tissue was homogenized using 0.5 mm glass beads with a TissueLyser LT homogenizer (Qiagen, Germantown, MD) to create a fine powder. DNA was extracted from the homogenized tissue using the E.Z.N.A Fungal DNA Extraction Kit (Omega Bio-tek, Norcross, GA). The total quantity of DNA extracted from each sample was quantified using a NanoDrop™ One Microvolume UV-Vis Spectrophotometer (Thermo Fisher Scientific, Rockford, IL). Fungal DNA in each sample was quantified using qPCR. A standard curve was created using *A. fumigatus* genomic DNA and consisted of 10-fold serial dilutions from 1 pg to 100 ng. 500 ng of sample DNA was analyzed per reaction and each sample was analyzed in triplicate. The sequences for the amplification primers and dual-labeled fluorogenic hybridization probe for the *A. fumigatus* 18S rRNA gene were previously published(8). All three probes were synthesized by Integrated DNA Technologies (IDT, Research Triangle Park, NC). Each 20 μL qPCR contained TaqMan® Fast Advanced Master Mix (Thermo Fisher Scientific, Rockland, IL), 500 ng of sample DNA and 1 μM each of forward primer, reverse primer and probe. Samples were run and analyzed using the Applied Biosystems™ 7500 Fast Real-Time PCR System (Thermo Fisher Scientific, Rockland, IL). The percent of fungal DNA in each sample was determined by dividing the quantified fungal DNA in each PCR by 500 ng (input DNA per sample). The total DNA obtained from each lung sample was multiplied by the percent fungal DNA to obtain the total fungal DNA/lung.

**Colony Forming Unit (CFU)**

Wildtype or Dectin-2$^{-/-}$ mice were infected with $4 \times 10^7$ conidia of *A. fumigatus* strain CEA10 through oropharyngeal inhalation under isoflurane anesthesia**.** Each experiment consisted of 5 wild-type and 5 Dectin-2$^{-/-}$ mice infected with CEA10 *Aspergillus* and one mock infection (PBS only) control per mouse line. After 48 hours lungs were extracted and placed into conical tubes containing sterile PBS. The weight of each tube pre- and post- lung addition was recorded to determine the weight of the tissue. The lungs were homogenized (Omni Tissue Homogenizer, Omni International, Kennesaw, GA) for 2 minutes each and then serial diultions were performed and plates onto SBD agar. Plates were incubated for 48 hours at 30$^0$C and colonies were counted. Data are reported at the number of colony forming units per gram of tissue. There was no growth from PBS only control mice.

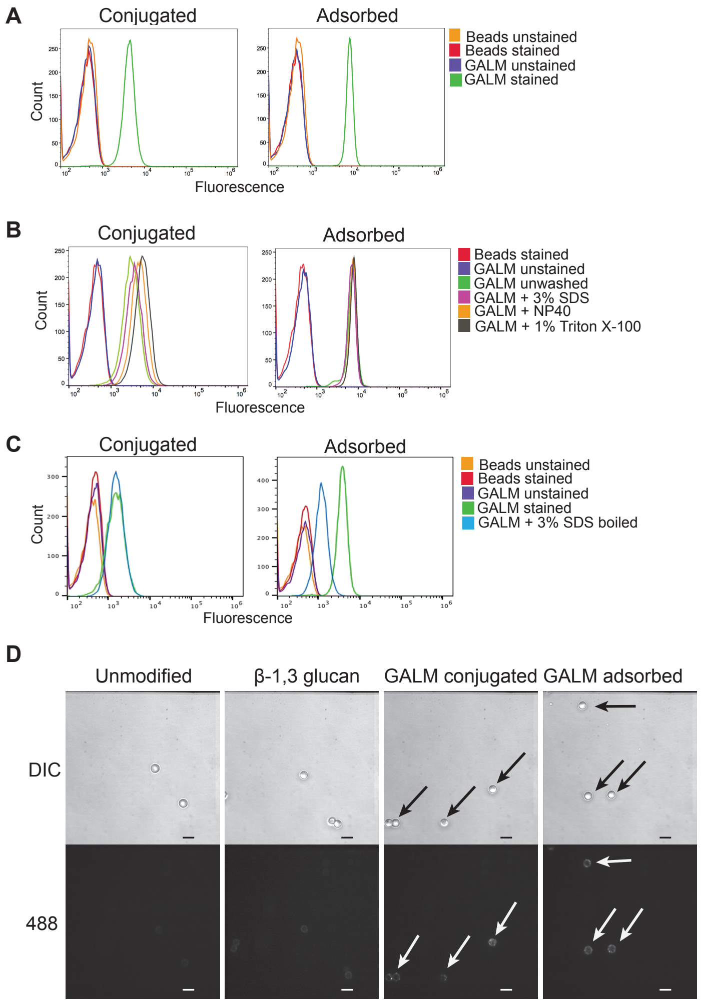


**Supplemental Figure S1**. Generation and validation of galactomannan FLP (A) Galactomannan FLP were created through adsorption or conjugation of purified *Aspergillus* galactomannan to the surface of amine-coated polystyrene beads. Flow cytometry comparing unmodified FLP and galactomannan FLP either unlabeled or incubated with anti-galactomannan antibody, followed by a secondary conjugated to Alexa Fluor 488 to detect galactomannan. Increased fluorescence indicates presence of galactomannan on the surface of the FLP. (B) Stability of carbohydrate attachment was assessed by incubating galactomannan FLP with common laboratory detergents for 1 hour, including Triton X-100, 3% SDS, and NP40. FLP were then washed and analyzed by flow cytometry. For both conjugated and adsorbed FLP, detergent washing did not disrupt surface association of the galactomannan to the FLP core. (C) FLP were then boiled for 1 hour in 3% SDS and assessed for carbohydrate stability. Under these stringent conditions, there was slight reduction in surface association of galactomannan from adsorbed FLP, but not conjugated. (D) Immunofluorescence of galactomannan FLP. Unmodified FLP,β-1,3 glucan FLP, and galactomannan FLP were labeled with anti-galactomannan antibody followed by secondary labelled with AF488 and analyzed using Spinning Disc confocal microscopy. Arrows indicating galactomannan FLP. Scale bar = 5 nm.

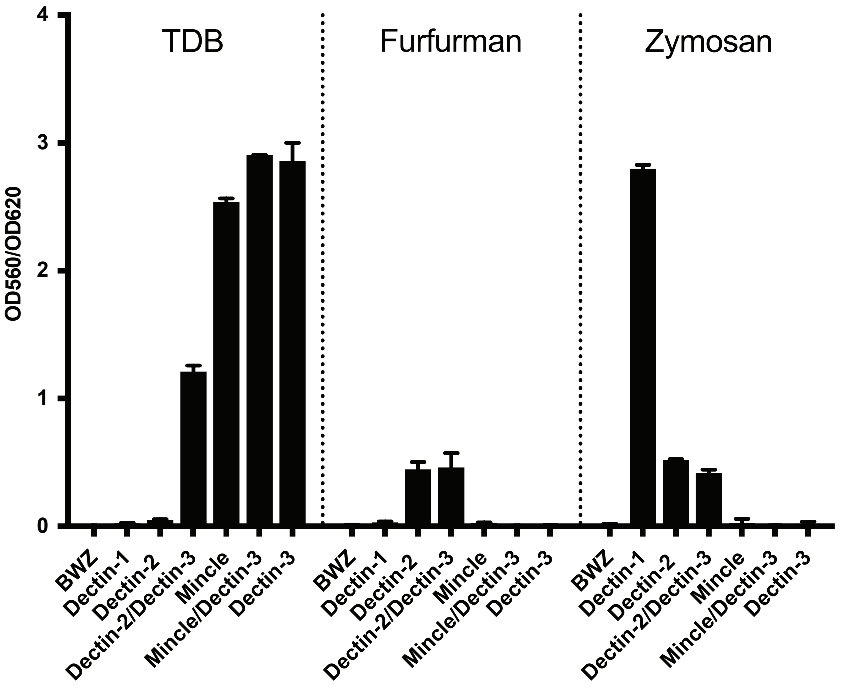


**Supplemental Figure S2.** CLR reporter cell activation using CLR control agonists. Reporter cell lines were stimulated for 18 hours with 10 µg/mL TDB (Mincle, Dectin-3 agonist), 10 µg/mL Furfurman (Dectin-2 agonist), or 100 µg/mL Zymosan (Dectin-1, Dectin-2 agonist). β-galactosidase activity was measured in total cell lysates using CPRG as a substrate. Error bars represent the SEM of duplicate wells, each experiment was repeated three times. As expected, TDB stimulated β-galactosidase production in cells containing either Mincle or Dectin-3, Furfurman stimulated Dectin-2 containing cells, and Zymosan stimulates Dectin-1 or Dectin-2 containing cell lines. Of note, the intrinsic β-galactosidase producing capacity of each cell line differs, therefore cross-cell line comparisons regarding the strength of the signal does not necessarily indicate differences in binding or signaling efficiency to a ligand.

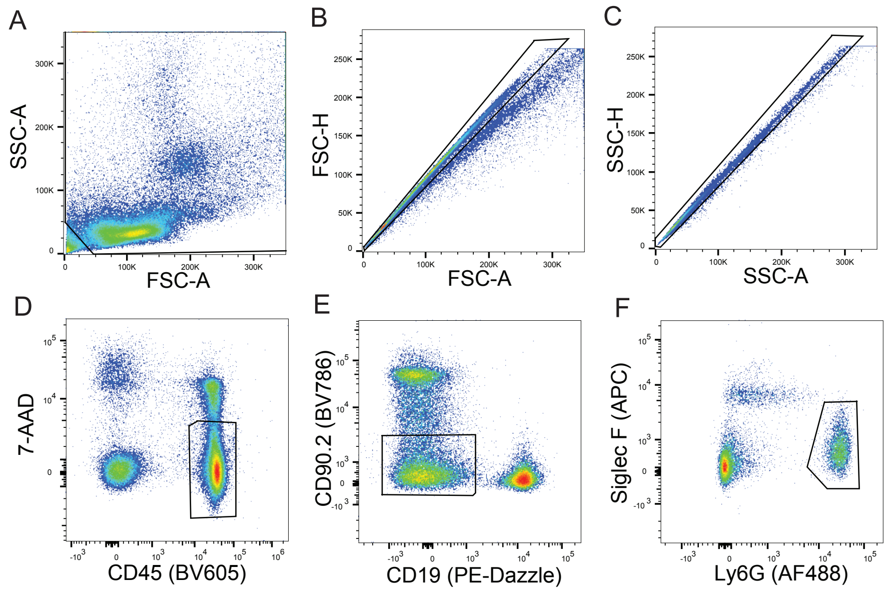


**Supplemental Figure S3.** Flow cytometry gating strategy for lung immunophenotyping to identify neutrophils. (A) Samples were first gated based upon FSC-A vs SSC-A to separate debris from cells (B and C) Cells were then gated using FSC-H vs FSC-A (B) followed by SSC-H vs SSC-A (C) to select for single cells. (D) Single cells were then gated using 7-AAD to identify live (7-AAD$^{lo}$) vs dead cells (7-AAD$^{high}$) and CD45-BV605 to identify live CD45$^{+}$ cells (7-AAD$^{lo}$ CD45$^{+}$). (E) The population of live CD45$^{+}$ cells were then separated based on CD90.2-BV786 and CD19-PE-Dazzle to separate T-cells (CD90.2$^{+}$ CD19$^{-}$) and B-cells (CD90.2$^{-}$ CD19$^{+}$) from innate immune cells (CD90.2$^{-}$ CD19$^{-}$). (F) The innate cell population was further separated using SiglecF-APC and Ly6G-AF488. Neutrophils were identified as SiglecF$^{lo}$ Ly6G$^{high}$ population.
